# Evaluating the Potential of Younger Cases and Older Controls Cohorts to Improve Discovery Power in Genome-wide Association Studies of Late-onset Diseases

**DOI:** 10.1101/693622

**Authors:** Roman Teo Oliynyk

## Abstract

For more than a decade, genome-wide association studies have been making steady progress in discovering the causal gene variants that contribute to late-onset human diseases. Polygenic late-onset diseases in an aging population display the risk allele frequency decrease at older ages, caused by individuals with higher polygenic risk scores becoming ill proportionately earlier and bringing about a change in the distribution of risk alleles between new cases and the as-yet-unaffected population. This phenomenon is most prominent for diseases characterized by high cumulative incidence and high heritability, examples of which include Alzheimer’s disease, coronary artery disease, cerebral stroke, and type 2 diabetes, while for late-onset diseases with relatively lower prevalence and heritability, exemplified by cancers, the effect is significantly lower. Computer simulations have determined that genome-wide association studies of the late-onset polygenic diseases showing high cumulative incidence together with high initial heritability will benefit from using the youngest possible age-matched cohorts. Moreover, rather than using age-matched cohorts, study cohorts combining the youngest possible cases with the oldest possible controls may significantly improve the discovery power of genome-wide association studies.

## 1. Introduction

With a growing fraction of the population reaching advanced age, late-onset diseases (LODs) have become the leading cause of mortality and morbidity [1]. Some LODs such as macular degeneration [2–4] are primarily caused by a single or a small number of high-effect variants. Each such disease is individually relatively rare in the population, and the mutations causing the majority of such diagnoses are known. The OMIM Gene Map Statistics [5] compendium lists thousands of such gene mutations.

The most common LODs are polygenic, also called complex diseases. They include heart disease, cancer, respiratory disease, stroke, and notably Alzheimer’s disease and other dementias [6]. The object of genome-wide association studies (GWASs) is to detect associations between genetic variants and traits in population cohorts, predict individuals’ LOD liability and, based on this knowledge, formulate preventive recommendations and treatments, with the ultimate goal of applying personalized medical interventions based on the genetic makeup of each unique individual [7]. With whole-genome sequencing becoming more accessible with every passing year, GWASs are being applied to all areas of genetics and medicine. Yet polygenic LODs remain resistant to the discovery of sufficient causal gene variants that would allow for accurate predictions of an individual’s disease risk [3,8,9]. GWASs can implicate only a subset of single nucleotide polymorphisms (SNPs) that can typically explain a fraction of the genetic heritability of a polygenic LOD [7], despite the fact that LODs with varied symptoms and phenotypes show high heritability in twin and familial studies [10–18].

Two complementary scenarios can explain LOD heritability, and both contribute to the so-called GWASs’ missing heritability problem [19–22]. The common low-effect-size allele hypothesis states that LODs are primarily caused by a combination of a large number of relatively common alleles of small effect [23]. GWASs have been able to discover only a small number of moderate-effect SNPs, but a large number of smaller effect SNPs remain below GWASs’ statistical discovery power. The rare high-effect-size allele hypothesis proposes that LODs are caused by a relatively small number of rare, moderate- or high-effect alleles with a frequency below 1% that likely segregate in various proportions into subpopulations or families [24,25] and are similarly problematic for GWASs’ discovery. Both scenarios can contribute to observational facts, but their relative weights vary depending on the genetic architecture of an LOD [26]. It has been determined [27,28] that common low-effect-size variants very likely explain the majority of heritability for most complex traits and LODs. This study primarily focuses on such diseases.

Recently, Warner and Valdes [29] stated that “one of the criticisms raised against genetic studies is that they are far removed from clinical practice.” Performing GWASs with ever-larger cohort sizes achieves better and more complete discovery for a variety of LODs and traits, yet larger patient cohorts are associated with practical, logistic, ethical and financial limitations, and research continues on developing statistical and procedural methods to improve discovery efficiency and sensitivity. Traditionally, GWASs recommend homogeneity of cohort participants. A common approach is to adjust for known covariates, including age, with the goal of correcting or averaging out biases [30]. Several studies caution about the appropriateness and scope of covariate adjustments [31,32]. Usually, the same age window is targeted, although it has been suggested [33] that individuals with an early age of onset are likely to have greater genetic susceptibility. Li and Meyre [33] proposed that once the risk of false positive association has been ruled out by initial replication studies, association can be extended to different age-matched windows. The recognition that “extreme phenotype sampling” may improve GWAS discovery prompted theoretical interest in study cohorts that are diverse in age [34,35].

A recent study [36] simulated population age progression under the assumption of relative disease liability remaining proportionate to individual polygenic risk, and determined that individuals with higher risk scores will become ill and be diagnosed proportionately earlier, bringing about a change in the distribution of risk alleles between new cases and the as-yet-unaffected population in every subsequent year of age. This is accompanied by lowering of the mean polygenic risk score (PRS) of the progressively older as-yet-unaffected population, and impairment of GWASs’ statistical discovery power for the study cohorts comprised of older age-matched individuals, most prominently for the highest prevalence LODs.

This research quantifies the use of non-age-matched cohorts for improving the discovery power of GWASs using as a case study eight prevalent LODs: Alzheimer’s disease (AD), type 2 diabetes (T2D), coronary artery disease (CAD), cerebral stroke and four late-onset cancers: breast, prostate, colorectal and lung cancer. The simulation results showed that GWASs of polygenic LODs that display both high cumulative incidence at older age and high initial familial heritability may benefit most from using the youngest possible participants as cases. Additional improvement in GWASs’ discovery power could be achieved by study cohorts that combine the youngest possible cases with the oldest possible controls.

## 2. Results

### 2.1. Impairment of GWASs’ statistical discovery power with progressively older age-matched cohorts

The preceding study [36] reported the patterns of GWASs’ discovery power for the age-matched cohorts. Out of the range of genetic architectures, simulation scenarios and validations performed in that study, it is necessary to refer to the findings for the common low-effect-size genetic architecture, which are summarized in this subsection for further comparisons.

The simulations in [36] determined that the age-related change in the cohort size needed to achieve 80% GWASs’ discovery power for an age-matched case-control cohort study increases with mid-cohort age (with the exception of lung cancer), as presented in Figure 1. This pattern is caused by the diminishing difference in effect SNP frequency between diagnosed cases and unaffected controls as mid-cohort age increases and is also reflected in the decreasing cohort PRS for older cohorts, see Figure S1 and Figure S2. This pattern was consistently observed for all genetic architectures, showing that the change in the PRS depends on the cumulative incidence and the magnitude of heritability. It is apparent that GWASs’ statistical discovery power diminishes as a complex function of a square of case-control allele frequency difference with cohort age according to Equations (6) and (7). Consequently, the age-matched cohorts composed of the youngest possible participants will allow for the best GWASs’ statistical discovery power compared to older age-matched cohorts.

**Figure 1.**
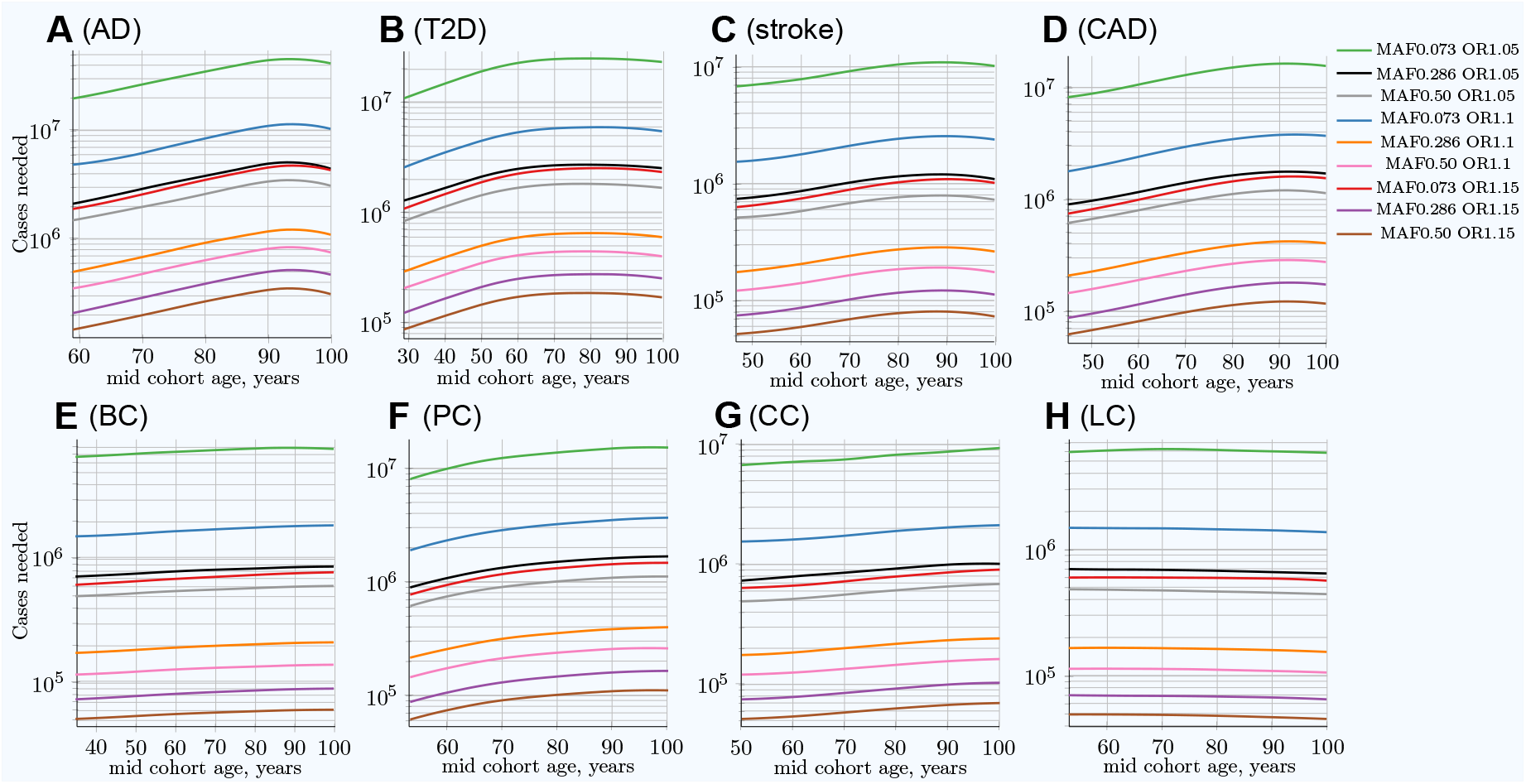
Change in number of cases needed to achieve 80% discovery power in age-matched cases and controls cohort design. **(A)** Alzheimer’s disease, **(B)** type 2 diabetes, **(C)** cerebral stroke, **(D)** coronary artery disease, **(E)** breast cancer, **(F)** prostate cancer, **(G)** colorectal cancer, **(H)** lung cancer. Age-matched cohorts require larger numbers of participants to achieve the same GWASs’ discovery power compared to the youngest cohort age. This figure was originally presented in [36].

The number of participants needed to achieve adequate GWASs’ statistical power differs between the lowest and the highest-effect alleles and also between the lowest and the highest frequency alleles, exhibiting a greater-than-hundredfold variation between alleles composing the genetic architecture, as seen in Figure 1. The required number of cohort participants is quite similar for the same-effect alleles among all eight LODs; for example, the highest-effect allele for each LOD requires 5· 10^4^−1.4· 10^5^ cases for 80% GWASs’ discovery power at younger ages. The change in allele frequency with age progression between cases and controls shows substantial variation among LODs, with the greatest change occurring in AD and the least significant in lung cancer, as demonstrated in Figure S1.

### 2.2. Advantage of using youngest possible cases and oldest controls in GWASs LOD cohorts

The scenarios simulating the number of cases needed when the cases are the youngest possible participants with increasingly older controls in the cohort are presented in Figure 2. In this scenario, the cohort size to achieve 80% GWASs’ statistical power decreases with the cohort age progression thanks to a change in allele frequency difference between younger cases and older controls cohorts; it is demonstrated in Figure S3. The multiplier representing the decrease in the number of cases that are needed in this scenario is represented by the blue lines in Figure 3, which strongly contrasts with the increasing with age multiplier of the number of cases needed for the same GWASs’ discovery power in the classic age-matched study design demonstrated by the red line. The age-matched and youngest case/older control scenarios are summarized in in Table 1. An additional side-by-side view can be seen in Figure S4. The youngest cases/older controls cohort scenario multiple was found to be almost identical between all allele frequencies and effect sizes for each particular LOD, as seen in Figure S5.

**Table 1.** Comparative summary of the older age-matched cohorts *and* the youngest cases-older controls cohorts *to* the youngest age-matched cohorts.

|  | Highly prevalent LODs |  |  |  | Cancers |  |  |  |
| --- | --- | --- | --- | --- | --- | --- | --- | --- |
|  | AD | T2D | Stroke | CAD | Breast | Prostate | Colorectal | Lung |
| LOD characteristics: |  |  |  |  |  |  |  |  |
| Lifetime risk % | 10–20 | 55 | 25–30 | 32–49 | 12 | 12 | < 4.5 | <6.9 |
| Heritability % | 79–80 | 69 | 38–44 | 50–60 | 31 | 57 | 40 | 8–18 |
| Maximum yearly incidence % | > 20 | 2.5 | 4.4 | 3.6 | <0.5 | <0.8 | <0.6 | <0.6 |
| Cohort size multiple for: |  |  |  |  |  |  |  |  |
| Age-matched at 80y | 1.82 | 2.13 | 1.51 | 1.86 | 1.15 | 1.65 (1.36) | 1.25 | 0.98 |
| Youngest cases & controls at 80y | 0.89 | 0.57 | 0.72 | 0.75 | 0.81 | 0.84 (0.82) | 0.90 | 0.88 |
| Relative advantage: 80y controls | 2.04 | 3.74 | 2.10 | 2.48 | 1.42 | 1.96 (1.66) | 1.39 | 1.11 |
| Cohort size multiple for: |  |  |  |  |  |  |  |  |
| Age-matched at 100y | 2.12 | 1.95 | 1.42 | 1.91 | 1.19 | 1.80 (1.44) | 1.36 | 0.92 |
| Youngest cases & controls at 100y | 0.43 | 0.46 | 0.46 | 0.52 | 0.72 | 0.72 (0.70) | 0.79 | 0.74 |
| Relative advantage: 100y controls | 4.39 | 4.24 | 3.09 | 3.67 | 1.65 | 2.50 (2.06) | 1.72 | 1.24 |

The MAF values and cases needed for 0.8 (80%) GWASs' statistical discovery power are for the common, low-effect-size alleles. Simulated cohorts span 10 years. The results shown are for the SNP with a MAF of 0.5 and an odds ratio (OR) of 1.15, the largest-effect allele, which requires the smallest cohort size for most accurate estimates. This effect is almost identical for the whole range of effect sizes and MAFs of the simulated SNPs within the genetic architecture. The two comparison blocks for 80-year and 100-year mid-cohort age compare the multiple of cases needed against the youngest possible age-matched cohorts for each LOD. The values show how many times larger (or smaller) cohort size the corresponding scenario would require to achieve the same 80% GWASs' statistical discovery power. Prostate cancer heritability is 57%, according to [37]. Shown in braces, 42% heritability [38], which, according to [36], is perhaps a more reasonable—although historically older—estimate of prostate cancer heritability and is more in line with the other three cancers.

**Figure 2.**
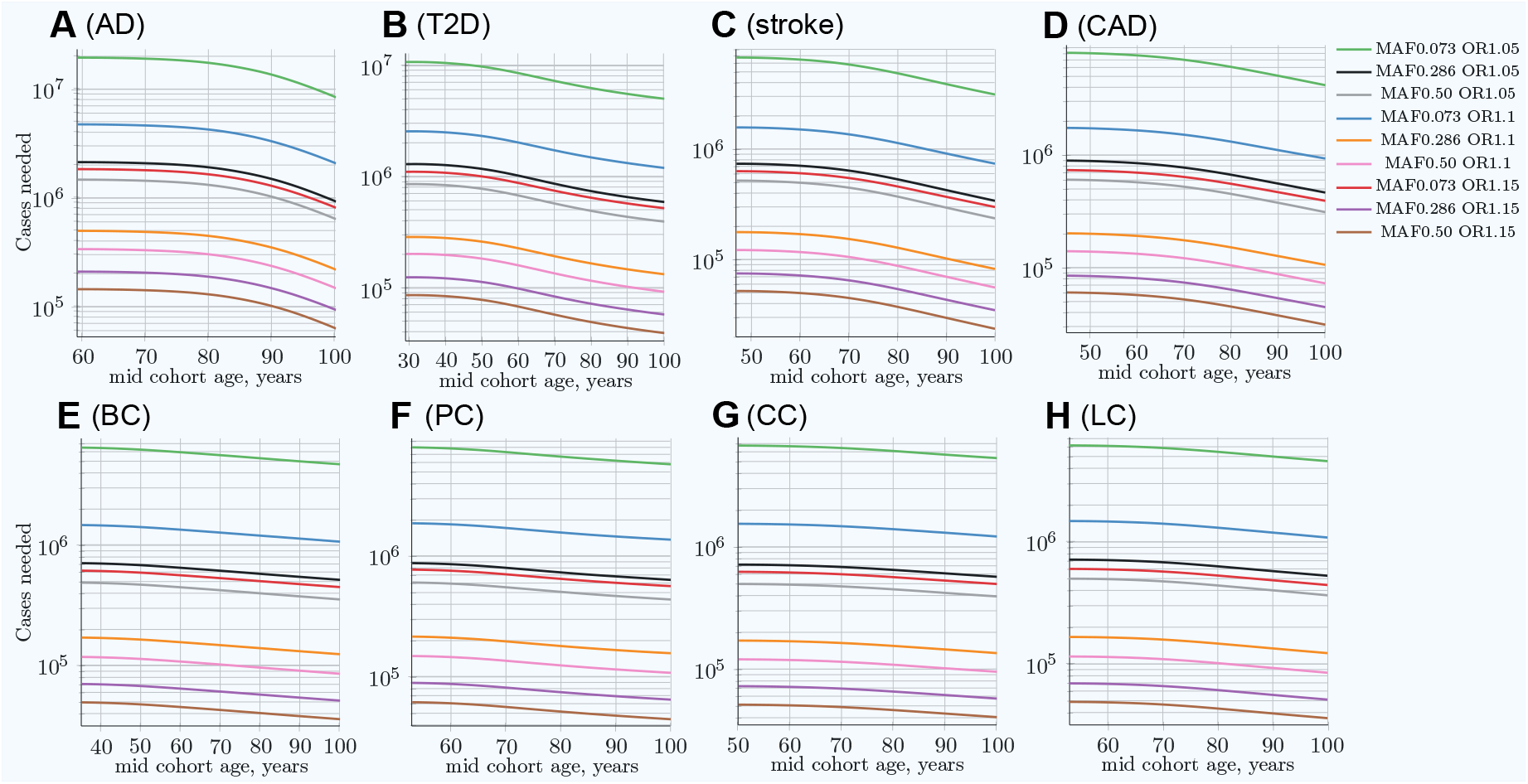
Change in number of cases needed for 80% discovery power in a cohort study when using progressively older controls compared to fixed-age young cases. **(A)** Alzheimer’s disease, **(B)** type 2 diabetes, **(C)** cerebral stroke, **(D)** coronary artery disease, **(E)** breast cancer, **(F)** prostate cancer, **(G)** colorectal cancer, **(H)** lung cancer. Cases’ mid-cohort age is leftmost age (youngest plot point); control mid-cohort ages are incremental ages. The number of cases needed for 80% discovery power is smaller when using older controls, particularly for those LODs showing the most prominent increase in the number of cases needed for older age in matched-age cohorts, as can be seen in Figure 1.

**Figure 3.**
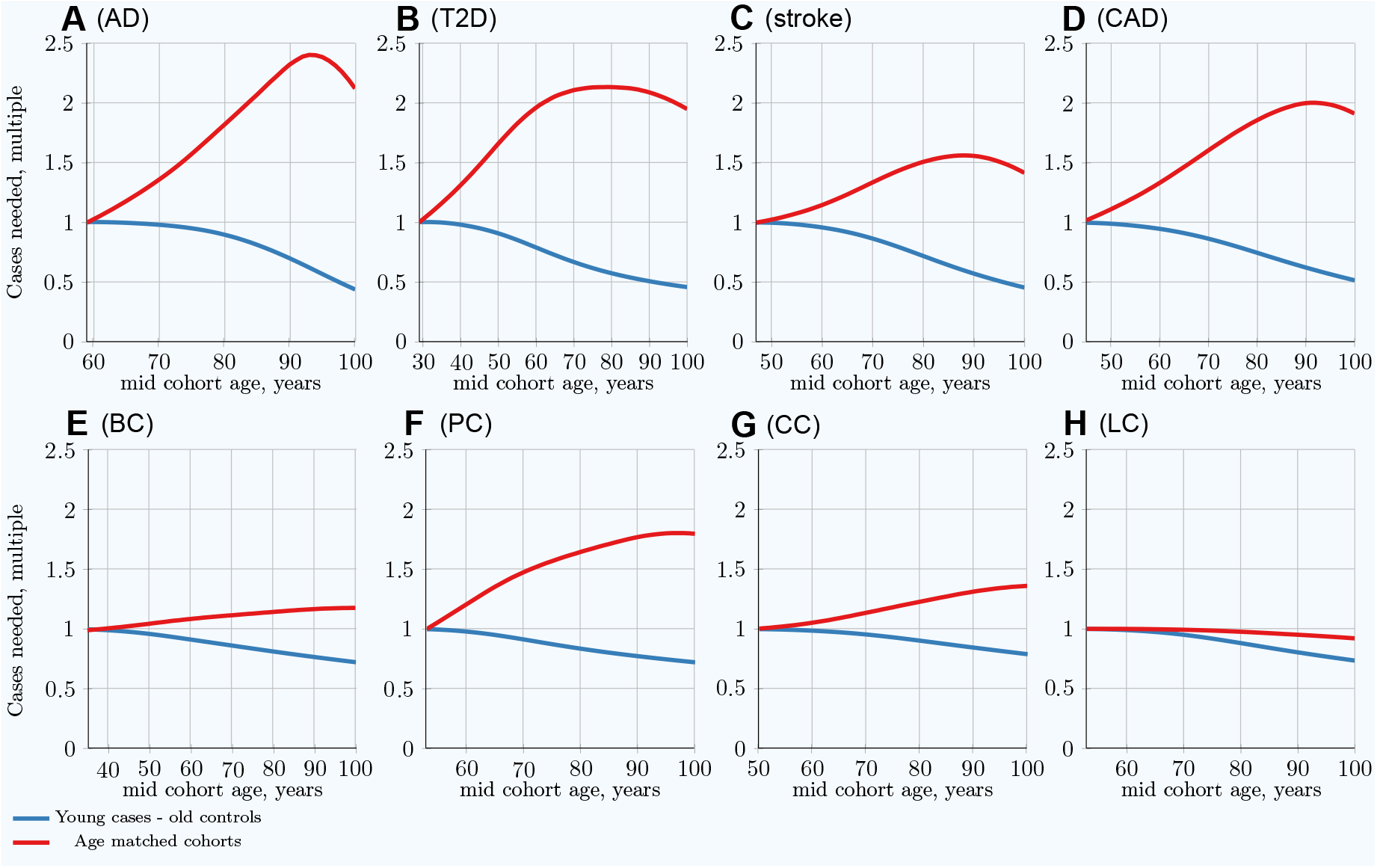
Per LOD comparison: Youngest possible cases and increasingly older controls *vs* classical age-matched cohorts. **(A)** Alzheimer’s disease, **(B)** type 2 diabetes, **(C)** cerebral stroke, **(D)** coronary artery disease, **(E)** breast cancer, **(F)** prostate cancer, **(G)** colorectal cancer, **(H)** lung cancer. The multiplier showing the reduction in the number of cases needed in a young cases-older controls scenario is shown in blue, strongly contrasting with the number of cases needed for the same GWASs’ discovery power in a classic age-matched study design, shown in red, which increases with age.

Thus, cohorts composed of the youngest possible cases and the oldest available controls could improve the discovery power of GWASs. Equivalently, such cohorts require even smaller numbers of participants to achieve the same GWASs’ discovery power than the youngest age-matched cohorts and certainly better than any older age-matched cohorts.

### 2.3. Characterizing and adjusting for effect size in the younger cases and older controls GWASs

The GWASs association analysis with the youngest possible cases and older controls cohorts showed that with the increasing controls cohort age, the SNP effect sizes exceed the known “true” effect sizes. This is the expected consequence of the larger effect allele differential between these cohorts compared with the age-matched cohorts. For SNPs defined in the simulation with the effect size 0.14 (OR=1.15), the association analysis found effect sizes near 0.20 (OR=1.21) for CAD and stroke with 100-year-old control cohorts; the bias is notably lower for the four cancers. The differential effect size (bias) of +0.05, corresponding to OR multiple equal 1.05, was reached for these LODs at control group age of 100 years; the bias age progression is displayed in Figure S6 and Figure S7. The typical single-SNP GWASs association analyses are known to show underestimated SNP effect for higher trait heritablities [39,40]. This is particularly relevant for AD and T2D with 3575 and 2125 of effect SNPs for common low-effect-size genetic architecture. Stringer *et al.* [39] consider this phenomenon a facet of GWASs’ missing heritability characteristic to single-SNP analysis. Multi-SNP analyses are proposed and are being developed [40–43]. For the purposes of this study, the customary single-SNP association analysis was sufficient for the relative bias determination for AD and T2D, which was found to closely follow the patterns of CAD and stroke, as well as the cancers.

The GWASs association analysis and effect-size adjustment with age and corresponding association standard errors are summarized in Table S1; the equations and approach are described in the Materials and Methods section 4.3. The progression shape in Figure S7 implies that the bias is proportionate to a power function by age, and the bias magnitude progression appears proportionate to the effect size magnitude. The linear regression fitted the normalized effect bias according to Equation (8) and Equation (9). The quadratic bias adjustment, used by Chatterjee *et al.* [44], resulted in a reasonable bias adjustment, as seen in Figure S8. The best fit power of age regression, with the power exponent specific for each LOD, produced a better match, although the slight improvement over the quadratic regression means that, for simplicity, the quadratic adjustment will be likely sufficient in practical GWASs bias correction for all LODs analyzed here (compare Figure S8 and Figure S9).

## 3. Discussion

By simulating population age progression under the assumption of relative disease liability remaining proportionate to individual polygenic risk, it was confirmed that individuals with higher risk scores will become ill and be diagnosed proportionately earlier, bringing about a change in the distribution of risk alleles between new cases and the as-yet-unaffected population in every subsequent year of age. With advancing age, the mean polygenic risk of the yet-unaffected aging population diminishes. The fraction of highest-risk individuals diminishes even faster, while at the same time, the LOD incidence increases or remains high with progression of age due to organism aging and cumulative environmental effects. Ultimately, the allele distribution in the as-yet-unaffected population of the same age with a given initial genetic architecture depends solely on cumulative incidence, which represents the fraction of the population that have succumbed to a disease [36]. GWASs’ statistical discovery power is impaired by the change in individual distribution of the PRS at older ages. A larger number of cases and controls is needed at older mid-cohort ages to achieve the same GWASs’ statistical discovery power compared to using younger age-matched cohorts. The effect is most prominent for AD, T2D, stroke and CAD, which exhibit higher heritability and cumulative incidence compared to the cancers analyzed here. The cancers show a noticeably smaller increase in the number of participants required to achieve the same statistical power, and while other factors could be at play, the probabilistic effects determined by lower incidence and lower heritability of the analyzed most prevalent cancers are sufficient to explain this pattern [36]. Quantitatively, the age-matched cohort studies would require 1.5-2.1 times more participants at age 80 compared to the youngest possible age-matched cohorts in the case of stroke, CAD, AD and T2D.

Designing cohorts composed of the youngest possible cases and the oldest available controls improves GWASs’ discovery power due to a larger difference in risk allele frequency between cases and controls. This effect is reminiscent of the example given by Sham and Purcell [45] for quantitative traits, in which performing GWASs using only the extreme top and bottom 5% of the individual distribution would achieve the same result with 4.4 times fewer participants compared to a cohort of randomly selected individuals. However, in contrast to Sham and Purcell [45] example, the observed larger MAF difference here is achieved not because of enrichment of effect alleles with age—the youngest case-control cohorts show the largest MAF difference and GWASs’ discovery power for the age-matched cohorts—but rather, the MAF difference effect is enhanced by impoverishment of the increasingly older controls in polygenic risk and corresponding effect allele frequencies. This cohort design leads to a smaller number of participants being needed for GWASs, particularly when applied to the highest cumulative incidence and heritability LODs—so much so that about 50% fewer participants are required to achieve the same GWASs’ statistical power when control cohorts between 90 and 100 years of age are matched to the youngest case cohorts, with the reverse being the case with older age-matched cohorts. Also notably, 20-25% fewer participants are needed in this scenario to achieve the same statistical power in cancer GWASs, including even those focusing on lung cancer.

Use of non-age-matched cases and controls in GWASs cohorts, while improving the discovery power, may result in reporting higher than “true” association effect, as should be expected with the enhanced difference in the effect of SNP frequency between cases and controls, and would require appropriate adjustment, as demonstrated in the Results section 2.3. This study’s simulations imply that the adjustment may be simplified by the fact that the bias magnitude was found proportionate to the associated SNP effect size. Many GWASs association software packages offer automated covariate bias correction [46–50] and allow for additional scripting.

Not every GWAS will be able to find a sufficient number of youngest cases as this study used as the basis of comparisons. However, due to a close to exponential rise in the incidence rate with age for most LODs at initial onset ages [36], the case cohorts can be formed at a somewhat older age or wider cohort span with correspondingly somewhat smaller improvement in GWASs’ discovery power. For all LODs analyzed, a majority of population would remain disease-free at ages 80- and 90-years old, with sufficient survivorship to provide a large pool of older controls.

The results of this study are based on the idealized simulations assuming that the gene-environment interaction, including the organism deterioration caused by the aging process, is following Cox’s proportional hazards model. The population in these simulations is homogeneous in all respects, while a practical GWAS would always have a varying degree of population diversity and nonhomogeneity that must be accounted for and addressed in GWASs’ quality control and study design [51]. While these concerns apply equally to age-matched and youngest possible cases-older controls study design, the advantage of the latter scenario may be realized to a varying degree in the practical GWASs.

In conclusion, the simulation results showed that GWASs of the polygenic LODs that display both high cumulative incidence at older age and high initial familial heritability will benefit from using the youngest possible participants. Moreover, GWASs would benefit from using as controls participants who are as old as possible. This may allow for additional increase in statistical discovery power thanks to achieving greater difference in risk allele frequency between cases and controls.

## 4. Materials and Methods

### 4.1. The simulation design summary and conceptual foundations

This study’s simulations are an extension of the author’s earlier research [36] that focused on the allele frequency and GWASs’ statistical power change patterns in aging populations for the eight LODs that were further analyzed here: Alzheimer’s disease, type 2 diabetes, coronary artery disease, cerebral stroke and four late-onset cancers—breast, prostate, colorectal, and lung cancer. A brief summary that includes excerpts from the Methods section of the earlier publication describing the model genetic architectures, the LOD incidence models, the statistical foundations and the simulation overview are provided in this subsection. Please see the Methods section in [36] for a more complete treatment. Subsections 4.2 and 4.3 will describe the simulation design and analysis that was performed exclusively in this study.

According to Chatterjee *et al.* [52], the conditional age-specific incidence rate of the disease, *I*(*t*|*G*) that is defined as the probability of developing the disease at a particular age *t*, given that a subject has been disease-free until that age, can be modeled using Cox’s proportional hazards model [53]:

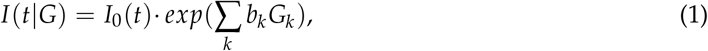

where *G* = (*G*_1_,…, *G*_*k*_) is the multiplicative effect of a set of risk factors on the baseline hazard of the disease *I*_0_(*t*). The set of age-independent variables in G could include genetic and environmental risk factors, as well as their interaction terms.

The following summary from Chatterjee *et al.* [52] is particularly relevant to the methodology of this research: “logistic regression methods are preferred for the evaluation of multiplicative interactions. For case-control studies, if it can be assumed that environmental risk factors are independent of the SNPs in the underlying population, then case-only and related methods can be used to increase the power of tests for gene-environment interactions. To date, post-GWAS epidemiological studies of gene-environment interactions have generally reported multiplicative joint associations between low-penetrant SNPs and environmental risk factors, with only a few exceptions.” This means that the polygenic score *G* = Σ_*k*_*b*_*k*_*G*_*k*_, as the lifelong characteristic of each individual, is used multiplicatively with *I*_0_(*t*), which encompasses environmental and aging effects. The simulations in this study used the functional approximations of the yearly incidence of Alzheimer’s disease, type 2 diabetes, coronary artery disease, and cerebral stroke and four late-onset cancers: breast, prostate, colorectal and lung cancer.

Five genetic architecture scenarios were analyzed in [36], and by comparing the patterns characteristic to each of these architectures, as well as extensive validation simulations, it was determined that the common low-effect genetic architecture, as indeed is the current scientific consensus [27,28], best fits the clinical and familial studies observations, and the analysis here is based exclusively on this architecture (although not discussed here, Supplementary Data also contains simulations and analysis results for rare medium-effect-size allele architecture).

In case of common low-effect genetic architecture, the MAFs are distributed in equal proportion at 0.073, 0.180, 0.286, 0.393 and 0.500, while the odds ratio (OR) values are 1.15, 1.125, 1.100, 1.075 and 1.05, resulting in 25 combinations. Having multiple well-defined alleles with the same parameters facilitated the tracking of their behaviors with age, LOD and simulation incidence progression.

An individual polygenic risk score *β* can be calculated as the sum of the effect sizes of all alleles, which is by definition a log(OR) (natural logarithm of odds ratio) for each allele, also following Pawitan *et al.* [54]:

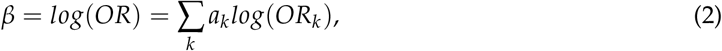

where *a*_*k*_ is the number of risk alleles (0, 1 or 2) and *OR*_*k*_ is the odds ratio of additional liability presented by the k-th allele. Variance of the allele distribution is determined by:

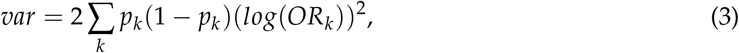

where *p*_*k*_ is the frequency of the k-th genotype [54]. The contribution of genetic variance to the risk of the disease is heritability:

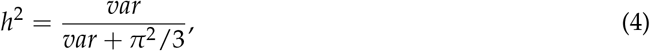

where *π*^2^/3 is the variance of the standard logistic distribution [55]. For example, the number of variants needed for the Scenario A LODs is summarized in Table 2.

**Table 2.**
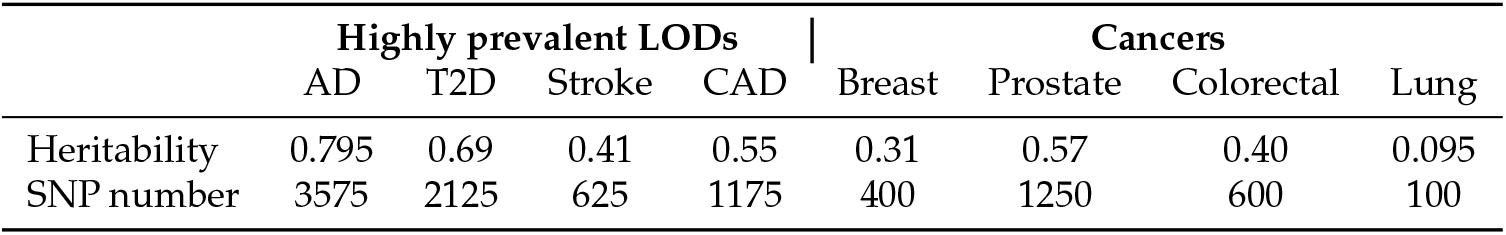
Heritability of analyzed LODs and an example of required variant numbers for common low-effect variants.

|  | Highly prevalent LODs |  |  |  | Cancers |  |  |  |
| --- | --- | --- | --- | --- | --- | --- | --- | --- |
|  | AD | T2D | Stroke | CAD | Breast | Prostate | Colorectal | Lung |
| Heritability | 0.795 | 0.69 | 0.41 | 0.55 | 0.31 | 0.57 | 0.40 | 0.095 |
| SNP number | 3575 | 2125 | 625 | 1175 | 400 | 1250 | 600 | 100 |

Following Pawitan *et al.* [54], the variants are assigned to individuals with frequencies proportionate to MAF *p*_*k*_ for SNP *k*, producing, in accordance with the Hardy-Weinberg principle, three genotypes (AA, AB or BB) for each SNP with frequencies 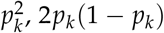 and (1 − *p*_*k*_)^2^. The mean value *β*_*mean*_ of the population distribution can be calculated using the following equation:

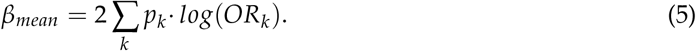

In this prospective simulation, each next individual to be diagnosed with an LOD is chosen proportionately to that individual’s relative PRS at birth relative to all other individuals in the as-yet-unaffected population. The number of individuals diagnosed annually is determined using the model incidence rate curve derived from clinical statistics. In this manner, the aging process is probabilistically reproduced using a population simulation model rather than a computational model. As the simulation progresses, the risk alleles are tracked for all newly diagnosed individuals and the remaining unaffected population, and their representation in the affected and remaining population is statistically analyzed. For each such allele in the simulated population, the allele frequency for cases and controls is tracked as age progresses. The difference between these MAFs gives the non-centrality parameter (NCP) *λ* for two genetic groups [45,56]:

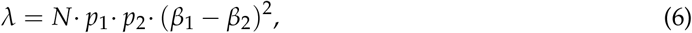

where N is the overall population sample size and *p*_1_*andp*_2_ the fractions of cases and controls, and *β*_1_ and *β*_2_ are the case and control mean effects for an allele of interest. The values *p*_1_ = *p*_2_ = 0.5, or an equal number of cases and controls, are used throughout this publication.

Having obtained NCP *λ* from Equation (6), Luan *et al.* [56] recommended using SAS or similar statistical software to calculate the statistical power, using the following SAS statement (an R equivalent statement was implemented in this study):

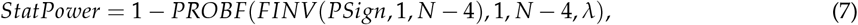

where *PSign* = 0.99999995 corresponds to the 5· 10^*−8*^ genome-wide significance level common in GWAS.

The comprehensive description of all simulation procedures and validation scenarios is available in [36].

### 4.2. Simulations and analysis of the youngest possible cases and older controls cohorts scenario

For the purposes of this research, rather than analyzing only the age-matched cohorts, the simulation was progressing in age until the mid-cohort age at which the fraction of population that succumbed to an LOD exceeded 0.25% population prevalence—the prevalence that was postulated as a minimum needed for forming the cases cohorts. This set of diagnosed individuals was kept for the duration of the simulation as the cases cohort. The cohort’s age span was fixed at 10 years just as in the preceding study, a relatively common cohort age span in GWASs. The simulation continued with population aging and being subject to probabilistic disease incidence, and at each progressive year of age, a new random set of as-yet-unaffected individuals was sampled, thereby forming a new cohort with progressively higher mid-cohort age. These cases and controls cohorts were analyzed for the effect allele frequency difference between cases and controls, with corresponding estimate of the cohort size needed to achieve 80% GWASs statistical discovery power. After completion of each simulation run with mid-cohort age exceeding 100 years, the results were aggregated and further analyzed.

### 4.3. GWASs association analysis and effect-size adjustment for younger cases and older controls cohorts

The case-control populations produced by these simulations were suitable for consequential GWASs association analysis that was implemented in this research. The simulations described in section 4.1 were extended to save the output in PLINK format [50,57]. The initial validation, analysis and file format conversions were performed using PLINK v1.9. The logistic regression GWASs with adjustment for age was performed using R script *AdjustByAge.R*, as described below, and the outputs were validated with Regression Modeling Strategies (rms) GWASs R package by Harrell Jr [49] and PLINK, confirming that the individual SNP association results with these two programs were identical to those produced by this R script.

The GWASs simulations showed that the apparent effect size tended to increase with age of control cohort, when analyzed against the youngest possible case cohort, compared to a “true” value, which was chosen as the effect size value from the youngest age-matched cohort. An example, although with a different objective, was demonstrated by application of age bias in a leprosy case-control study [44] that used the bias adjustment as a function of squared age.

The R script written for this analysis *AdjustByAge.R*, based on R Generalized Linear Models glm() functionality [58], performed the GWASs association and iterative age covariate adjustment starting with a youngest possible age-matched cohort and proceeding with the progressively older control cohorts. The script effectively discovered the best match bias adjustment power and allowed comparing the power parameters analyzed between LODs, as presented in the Results section. Importantly, the bias adjustment results showed that the increase in the value of the effect size was approximately proportionate to the effect size magnitude for all LODs analyzed here. The differential normalized effect size *D*(*t*) can be expressed as:

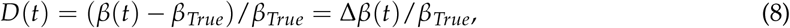

where *β*(*t*) and *β*_*True*_ are the effect-size values found for older control cohorts compared to a known “true” effect sizes as defined in the simulated genetic architecture for each allele. The variable *D*(*t*) will be referred to further as normalized bias. The GWASs simulations associated the effect sizes in 5-year control cohort age increments and matched the best power exponent regression function:

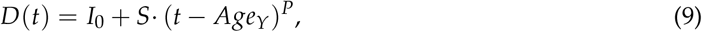

where *I*_0_ and *S* are the linear regression intercept and slope, *t* is an older control cohort age, *Age*_*γ*_ is the youngest case cohort age and *P* is the best match power exponent. When the solution to Equation (9) is correctly estimated for one gene variant (likely for an SNP with a larger effect size), it could be used to adjust other discovered variants’ effect sizes from Equation (8) and Equation (9). The R script *FindAdjustmentRegressionFunction.R* implementing lm() linear regression iteratively fitted the best matching adjustment power with lowest residuals. Additionally, this script evaluated the regression with fixed P=2 (quadratic regression) for all LODs. The data preparation and scripting steps described here are listed with specifics in *GwasSimulationPipeline.txt*, available along with the R scripts in Supplementary Data.

### 4.4. Data sources, programming and equipment

The population mortality statistics from the US Social Security Actuarial Life Table [59] provided yearly death probability and survivor numbers up to 119 years of age for both men and women. Disease incidence data from the following sources were extensively used for analysis, using the materials referenced in supplementary Chapter S1 in [36]: Alzheimer’s disease: [11,60–62]; type 2 diabetes: [63]; coronary artery disease and cerebral stroke: [64]; and cancers: [65,66].

The simulations were performed on an Intel Xeon Gold 6154 CPU-based 36-core computer system with 288GB of RAM. The simulation is written in C++ and can be found in Supplementary Data. The simulations used population pools of 2 billion individuals for the LOD simulations and 300 million for validation simulations, resulting in minimal variability in the results between runs. The cohort simulations were built sampling at minimum 5 million cases and 5 million controls from the surviving portion of the initial 2 billion simulated individuals, which is equivalent to 0.25% of the initial population. This means that the cohort study began its analysis only when this cumulative incidence was reached. Conversely, the analysis ceased when, due to mortality, the number of available cases or controls declined below this threshold. For all LODs, this maximum mid-cohort age was at least 100 years and, depending on LOD, up to a few years higher. This confirms that, as described later in the Discussion section, in cohorts composed of younger cases and older controls, it is feasible to form control cohorts of up to 100 years of age.

The simulation runs for either all validation scenarios or for a single scenario for all eight LODs took between 12 and 24 hours to complete. The final simulation data, additional plots and elucidation, source code and the Windows executable are included in Supporting Information. Intel Parallel Studio XE was used for multi-threading support and Boost C++ library for faster statistical functions; the executable may be built and can function without these two libraries, with a corresponding slowdown in execution. The ongoing simulation results were saved in comma separated files and further processed with R scripts during subsequent analysis, also available in Supplementary Data.

### 4.5. Statistical analysis

Large variations between simulation runs complicate the analysis of population and genome models. This issue was addressed in this study by using a large test population, resulting in negligible variability between runs. The statistical power estimates deviated less than 1% in a two-sigma (95%) confidence interval, except for the early Alzheimer’s disease cohort, which commenced at 1.5% and fell below the 1% threshold within 4 years (see \**TwoSDFraction.csv* files in Supplementary Data). In addition to ensuring that the simulations operated with reliable data, this eliminated the need for the confidence intervals in the graphical display.

The GWASs simulations and variant effect size covariate adjustment by age were more memory-intensive, and the 200 million simulated population with 500 thousand case and control cohorts was possible with the described equipment. In this instance, two-sigma confidence intervals for simulated GWASs discovery and regression parameters are presented in the corresponding plots.

## Acknowledgments

The author thanks Alexei J. Drummond at the University of Auckland for a number of helpful and challenging discussions.

## Abbreviations

The following abbreviations are used in this manuscript:

AD: Alzheimer’s disease
CAD: coronary artery disease
GWAS: genome-wide association study
LOD: late-onset disease
MAF: minor allele frequency; customarily implying the ‘effect allele frequency’
OR: odds ratio
PRS: polygenic risk score
SNP: single nucleotide polymorphism; in context of this study used synonymously with the term ‘allele’
T2D: type 2 diabetes

## Appendix A. Supplementary Document

A document containing supplementary figures and tables is attached after the main manuscript.

## Appendix B. Supplementary Data

A zip file **SupplementaryData.ZIP** containing the simulation executable, the source code, R scripts, batch files, and simulation results.

## SUPPLEMENTARY FIGURES AND TABLE

**Figure S1.**
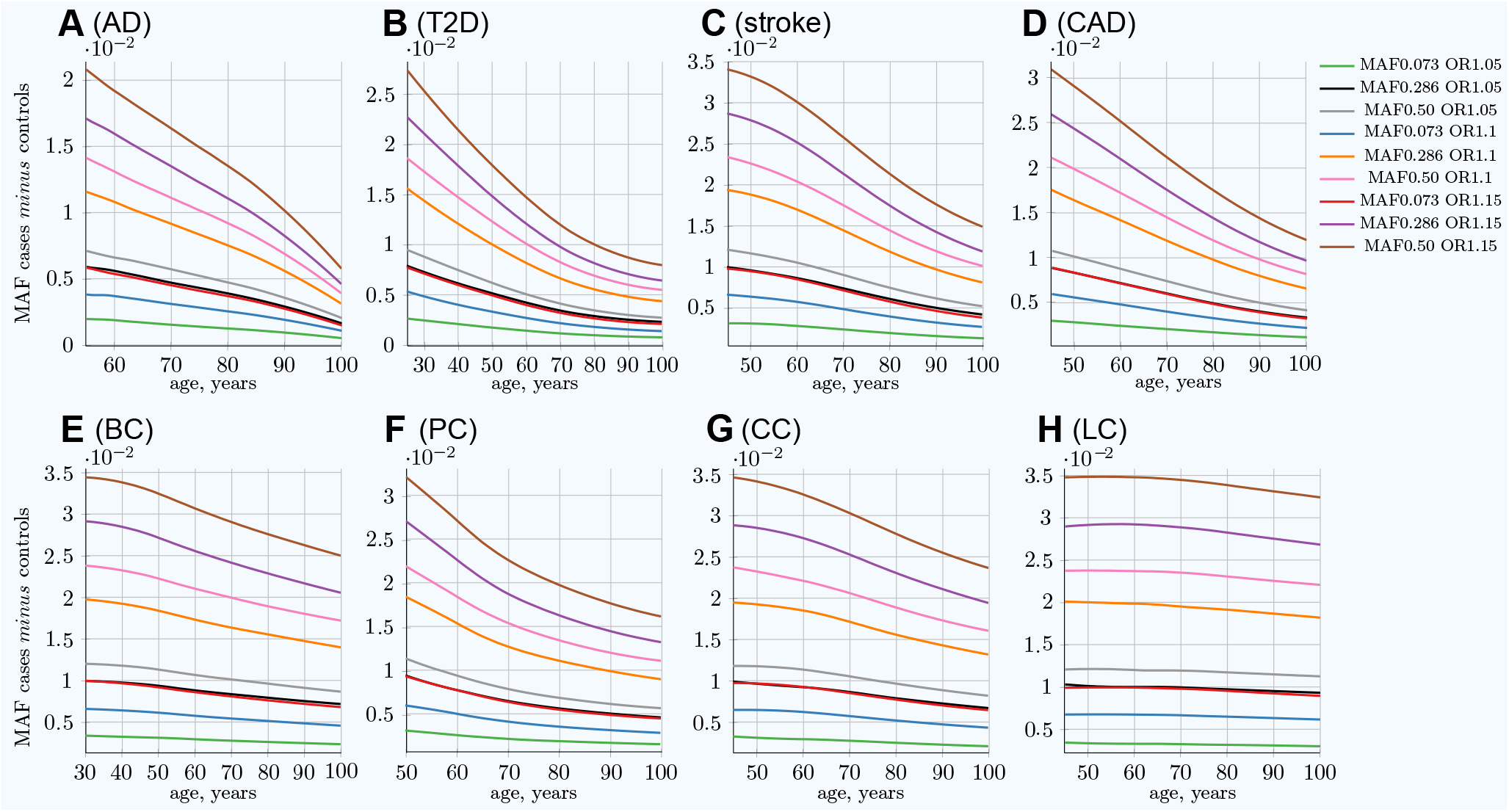
Difference in frequency in allele between newly diagnosed individuals and remaining population of the same age. **(A)** Alzheimer’s disease, **(B)** type 2 diabetes, **(C)** cerebral stroke, **(D)** coronary artery disease, **(E)** breast cancer, **(F)** prostate cancer, **(G)** colorectal cancer, **(H)** lung cancer. The **MAF cases *minus* controls** value is used to determine GWASs’ statistical power. Rarer and lower-effect-size (OR) alleles are characterized by a lower MAF relative change. *(Displayed here: nine out of 25 SNPs for common low-effect-size genetic architecture).* From Oliynyk (2019).

**Figure S2.**
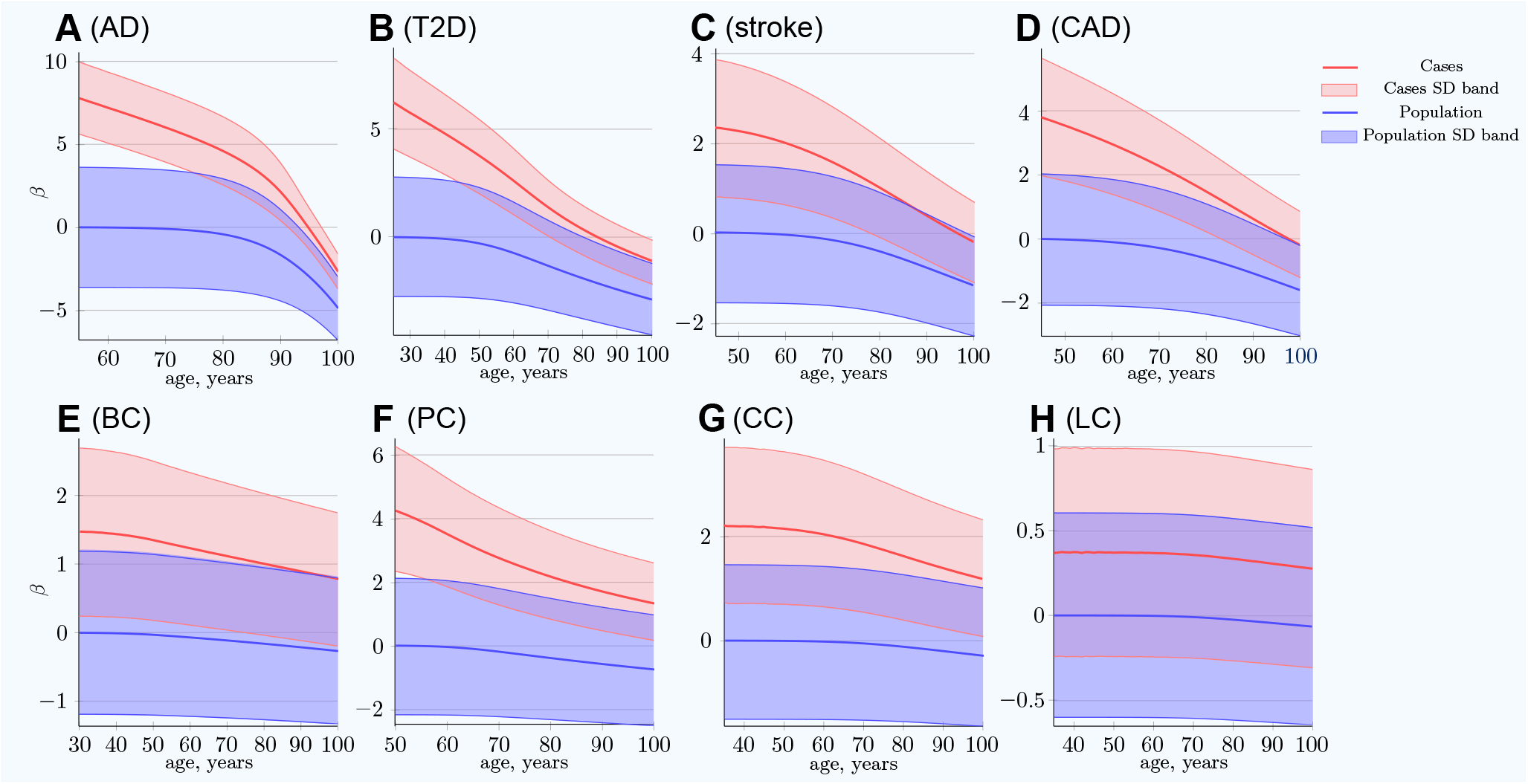
Polygenic risk score difference between newly diagnosed individuals and the remaining unaffected population. **(A)** Alzheimer’s disease, **(B)** type 2 diabetes, **(C)** cerebral stroke, **(D)** coronary artery disease, **(E)** breast cancer, **(F)** prostate cancer, **(G)** colorectal cancer, **(H)** lung cancer. *SD band* is a band of one standard deviation above and below the cases and the unaffected population of the same age. For highly prevalent LODs, at very old age, the mean polygenic risk of new cases crosses below the risk of an average healthy person at early onset age. *(Common low-effect-size alleles, showing largest-effect variant with MAF = 0.5, OR = 1.15).* From Oliynyk (2019).

**Figure S3.**
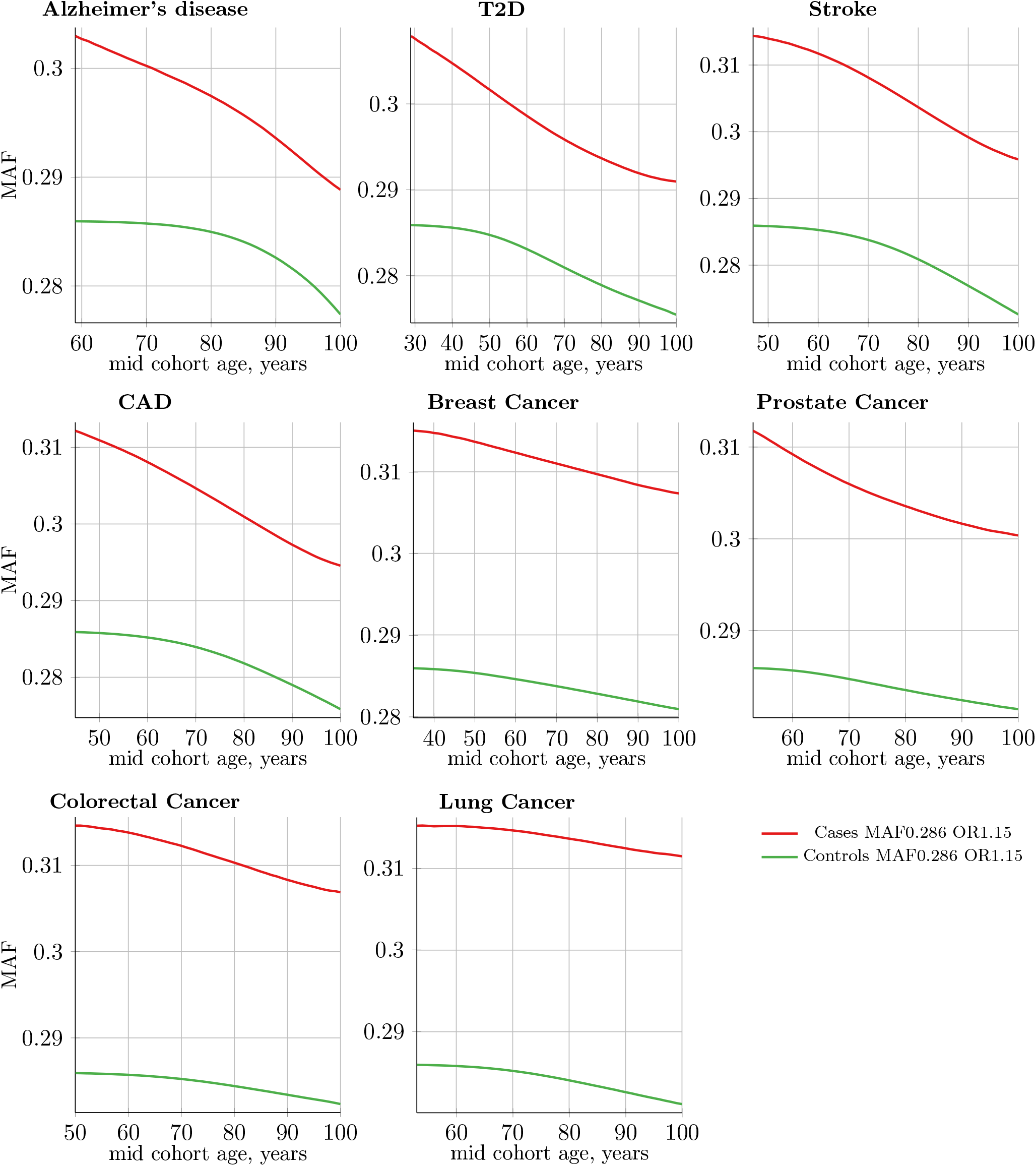
Absolute magnitude change in minor allele frequency (MAF) with age for cases and controls; cohort simulation. Common, low-effect-size alleles; all plots show MAF = 0.286 and OR = 1.15 allele. Change in the absolute magnitude of each allele frequency value is relatively small with age progression. GWASs’ discovery power is a function of the difference in allele frequency between cases and controls. It is easy to visually estimate the change in the difference in allele frequency between the cases and controls. In the age-matched scenario, the difference is taken between points on the line at the same mid-cohort age. For the youngest cases-older controls scenario, the difference is taken always between the leftmost point on the red line and progressively older controls on the green line. From Supplemental Information in Oliynyk (2019).

**Figure S4.**
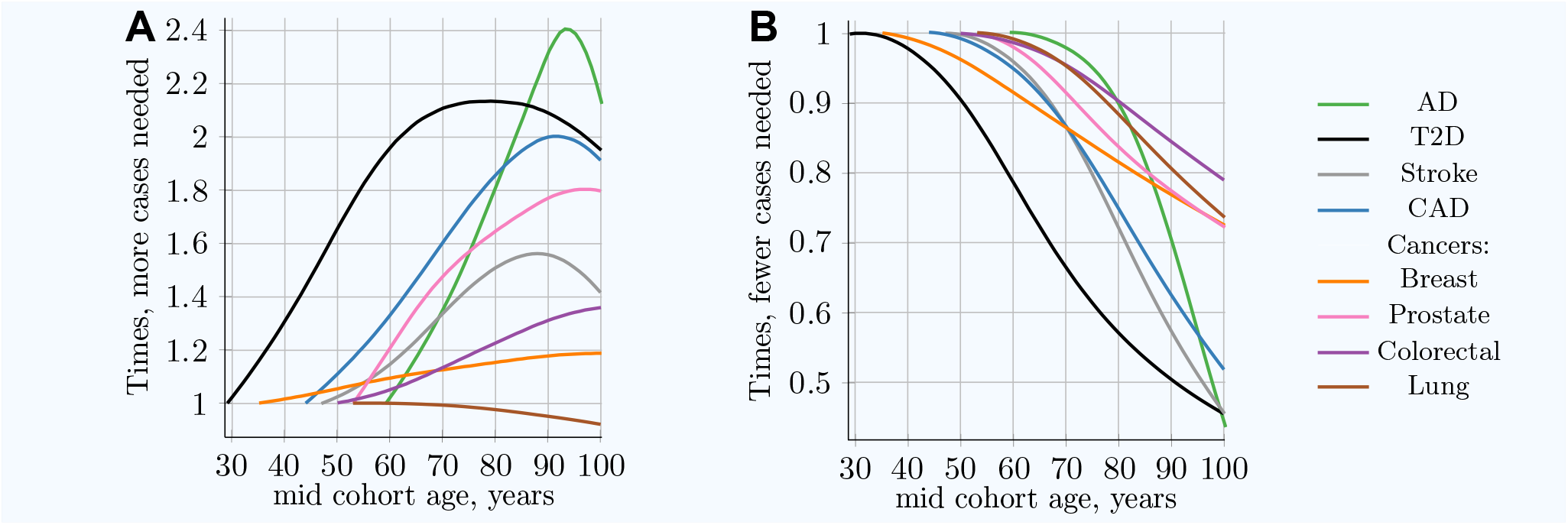
Advantage of using youngest possible cases and increasingly older controls compared to classical age-matched cohorts. **(A)** Relative **increase in number of cases needed** for 80% discovery power in a cohort study using progressively older **case and control cohorts of the same age. (B)** Relative **decrease in the number of cases needed** for 80% discovery power in a cohort study using **progressively older control cohorts compared to fixed-age young-case cohorts**. The youngest age cohort for each LOD is defined as the mid-cohort age at which the cumulative incidence for a cohort first reaches 0.25% of the population. Therefore, the leftmost point on each LOD line is the reference (youngest) cohort, and as cohorts age, the cohort case number multiple required to achieve 0.8 statistical power is relative to this earliest cohort. While all alleles display a different magnitude of cases needed to achieve the required statistical power, the change in the multiplier with age is almost identical for all alleles within a given genetic architecture scenario. *(Common low-effect-size genetic architecture)* Plot A of this figure from Oliynyk (2019).

**Figure S5.**
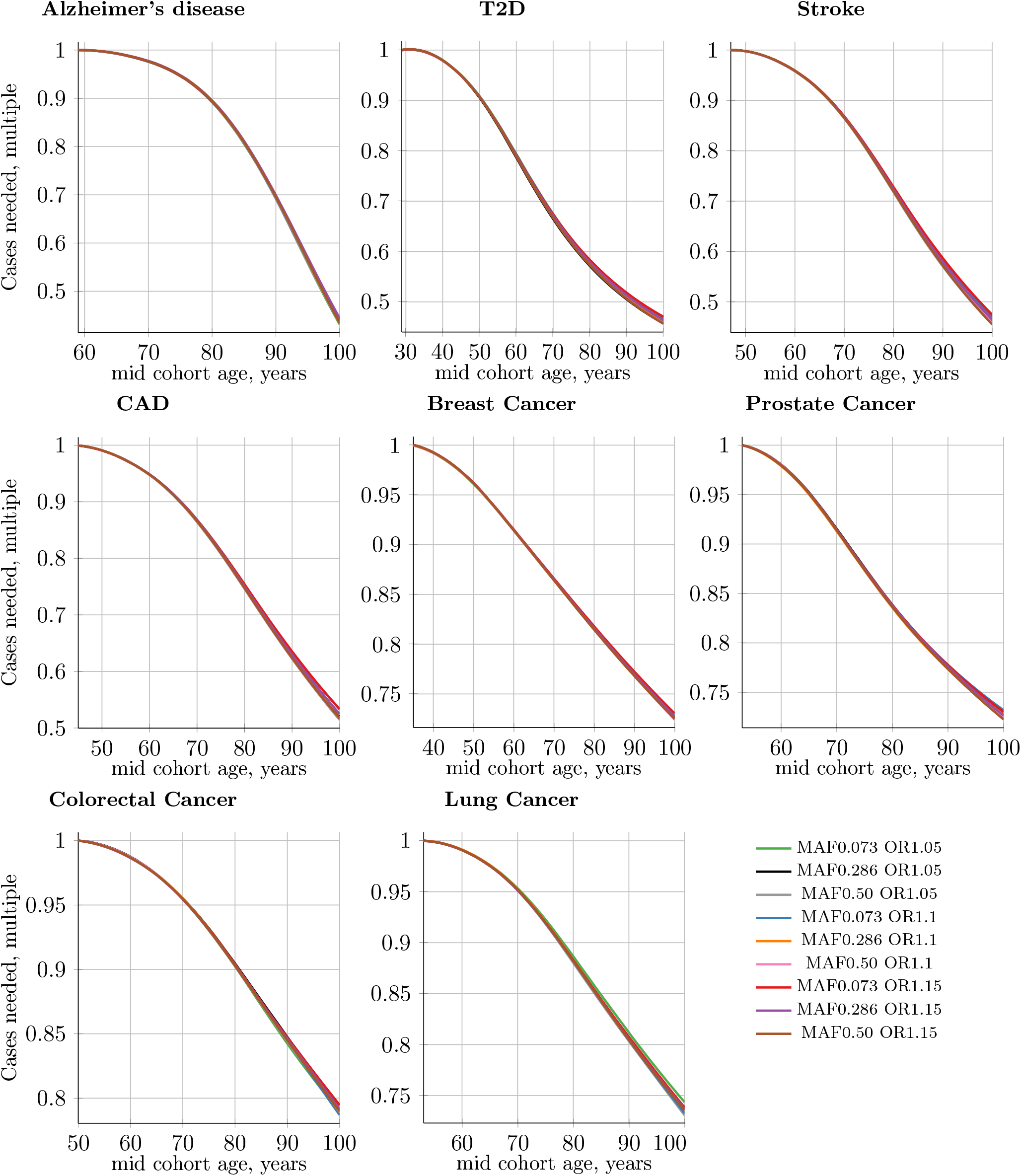
Multiple of the decline in the number of cases needed for 0.8 discovery power in a cohort study using progressively older control cohorts compared to a fixed-age young-cases cohort. Cases’ mid-cohort age is leftmost age (youngest plot point); control mid-cohort ages are incremental ages. The number of cases needed for 0.8 discovery power is smaller when older controls are used, particularly for LODs with the highest heritability and incidence. Common, low-effect-size alleles. A sample of nine out of 25 SNPs; MAF = minor (risk) allele frequency; OR = risk odds ratio.

**Figure S6.**
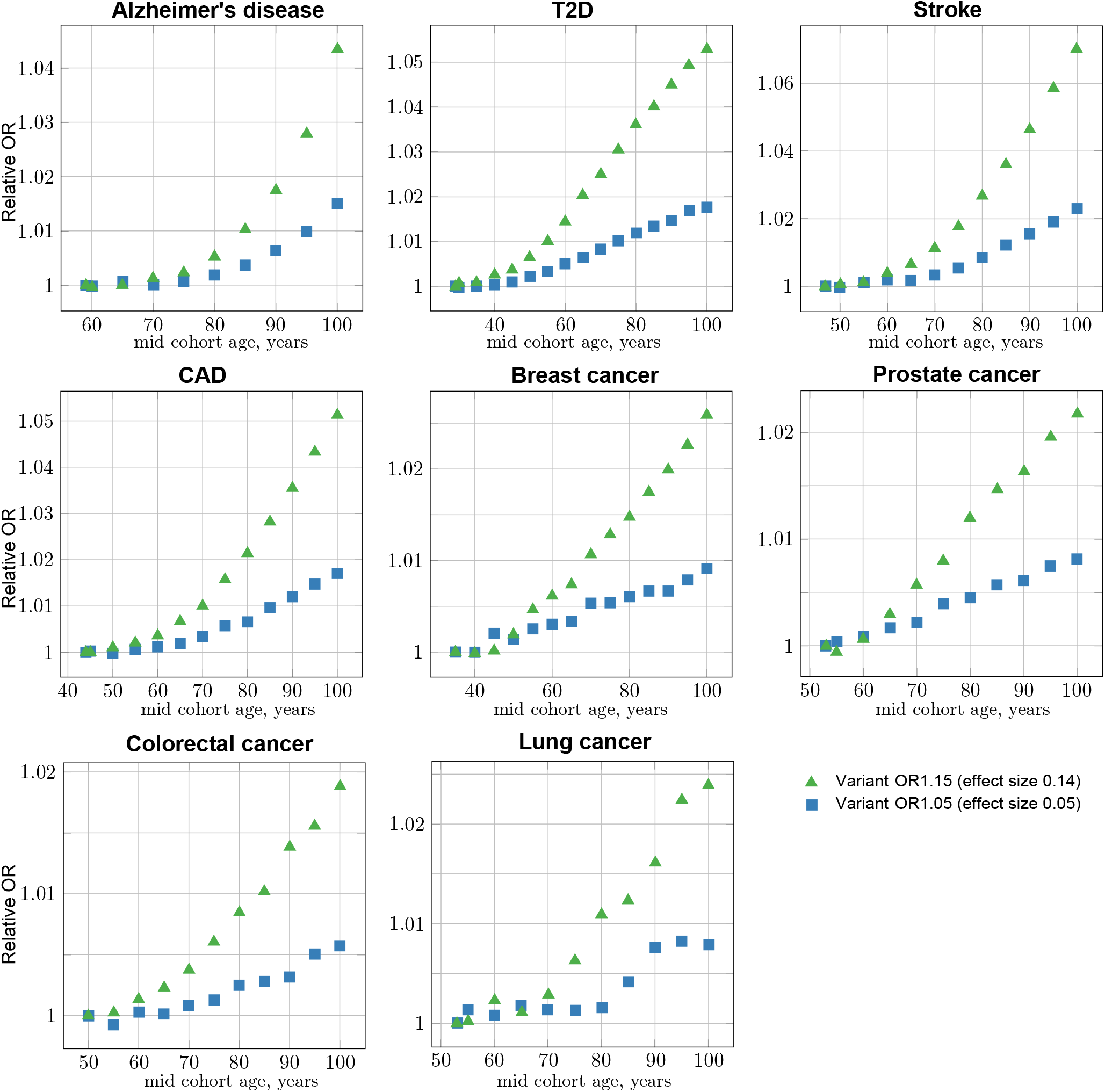
GWAS association simulations: OR bias progression with control cohort age increasing against the constant youngest possible case cohort. Common, low-effect-size alleles, showing two SNPs—with the largest and the smallest effect—for each LOD. The OR increase (bias) with mid-cohort age progression implies a power of ΔAge from age-matched youngest cohort. The confidence intervals are not displayed on this plot for illustration purposes; they are displayed in Figure S5, showing the same data in effect size units.

**Figure S7.**
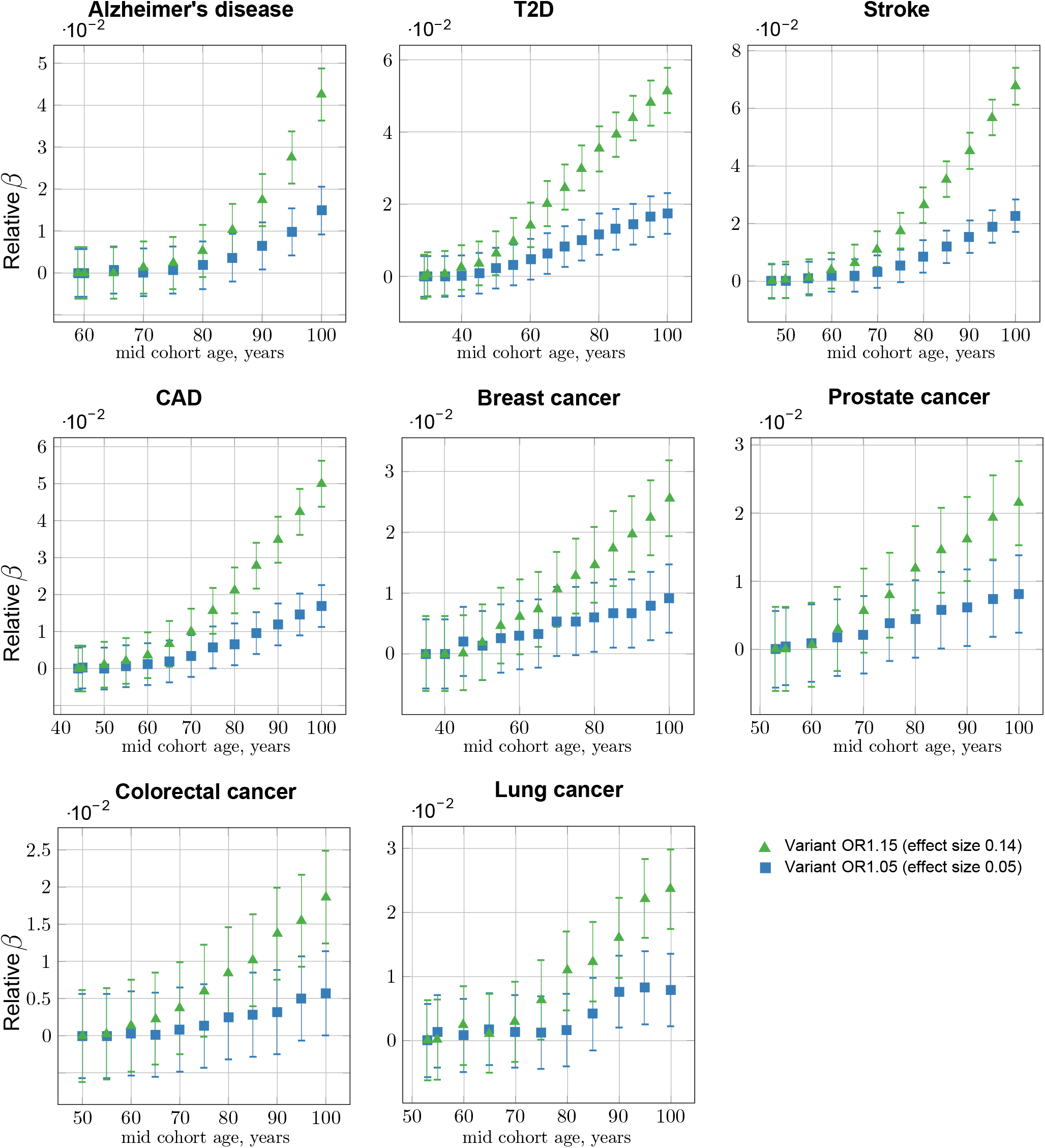
GWASs association simulations: relative effect size bias progression with control cohort age increasing against the constant youngest possible case cohort. Common, low-effect-size alleles, showing two SNPs—with the largest and the smallest effect—for each LOD. The confidence interval bars correspond to two-sigma (95%) confidence from the GWASs’ logistic regression association. The OR increase with mid-cohort age progression implies a power law relative to Δ*age*. This plot implies the LOD SNP age bias and corresponding adjustment value are proportionate to the SNP effect size; this observation is further investigated in Figure S6 and Figure S7.

**Figure S8.**
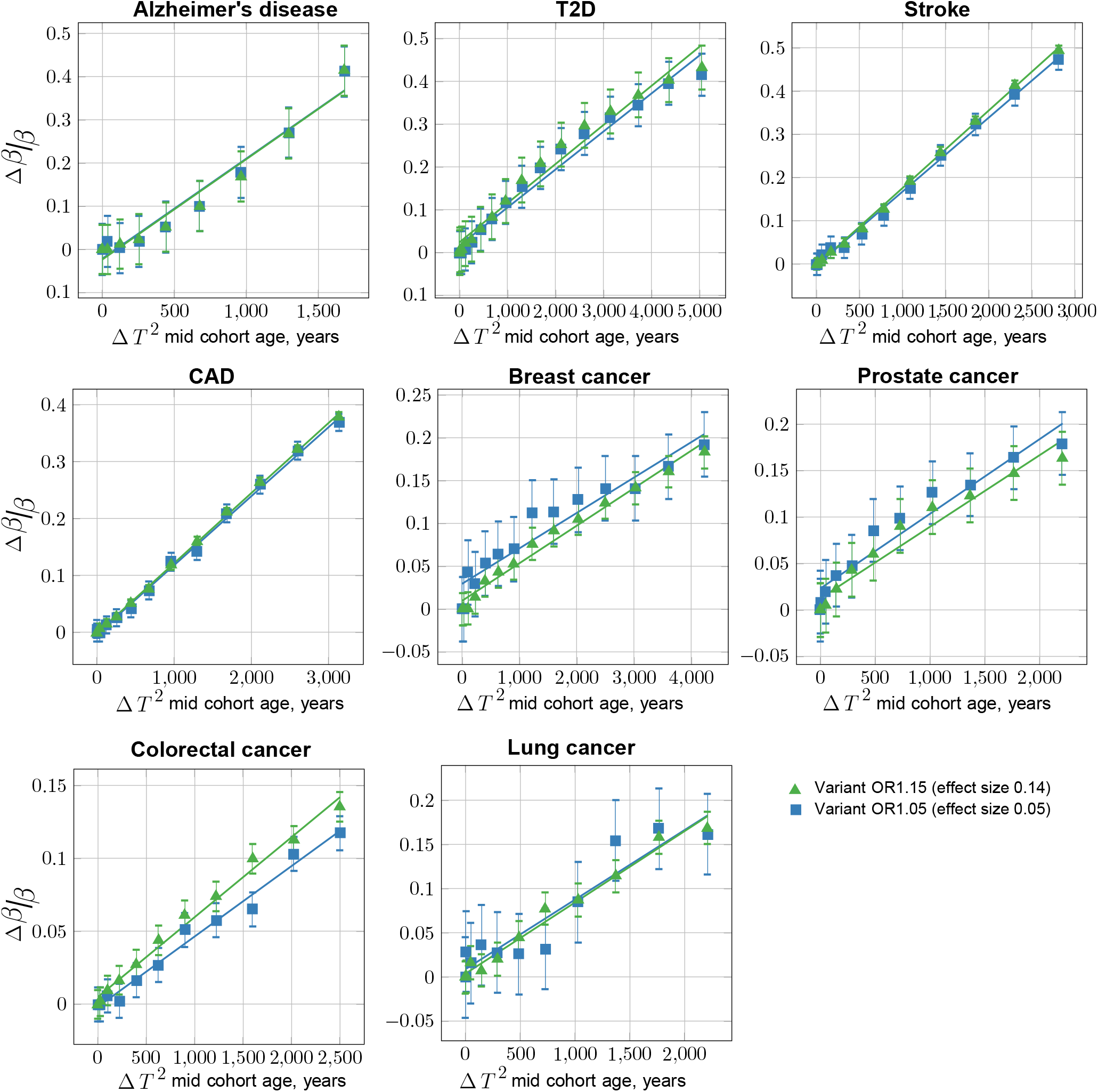
GWASs association simulations: characterizing the age bias adjustment maintaining “true” OR with control cohort age progression (quadratic: Δ*T*^2^). Common, low-effect-size alleles, showing two SNPs—with the largest and the smallest effect—for each LOD. The confidence interval bars correspond to two-sigma (95%) based on standard error of linear regression fitting. This plot depicts the adjustment proportionate to square of Δ*t* = *t* − *T*_*Y*_ – relative age from the youngest cohort mid-cohort age for the normalized bias of the effect size *β* calculated Δ*β/β*, as described in the main article. See also Table S1.

**Figure S9.**
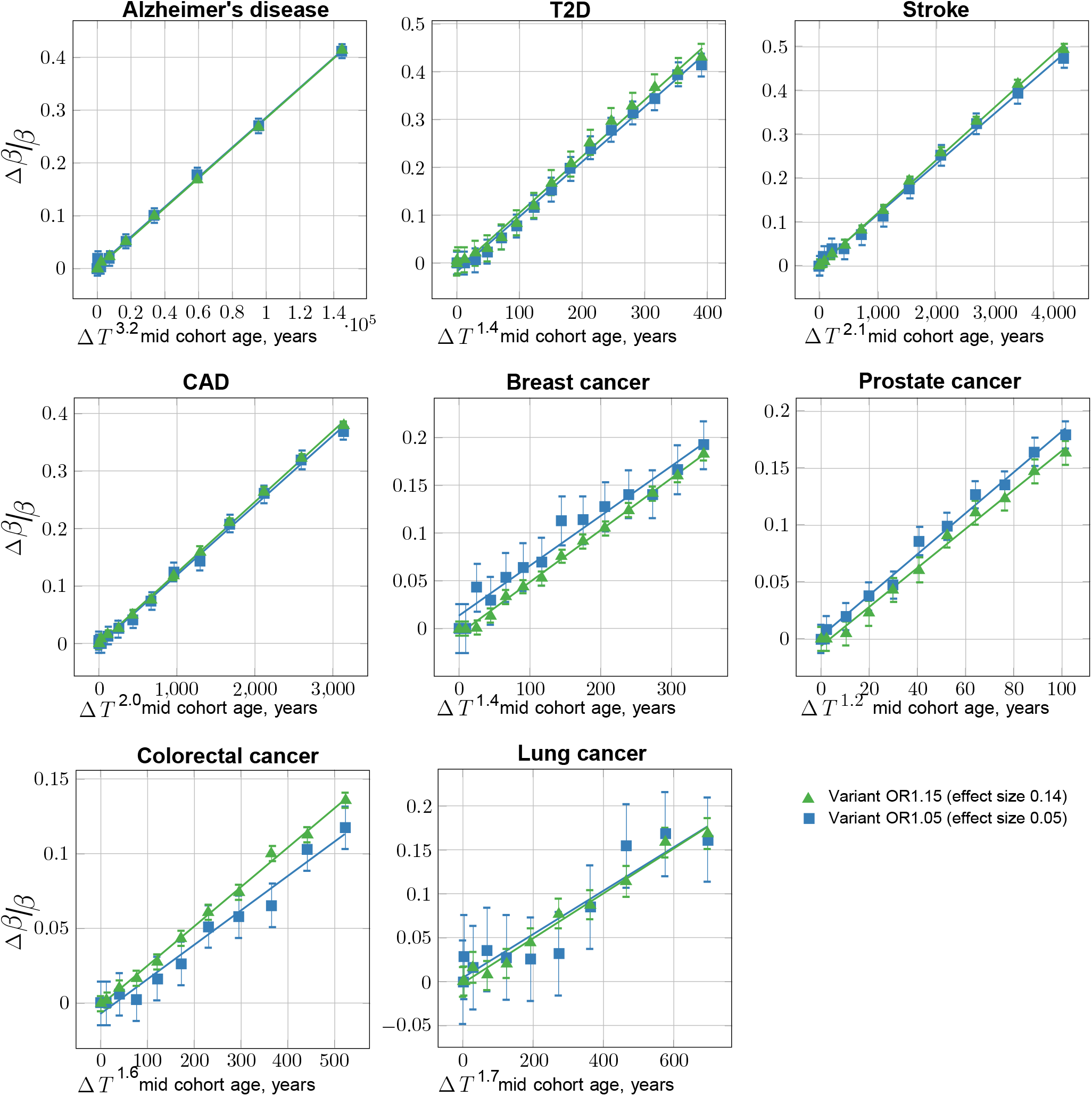
GWASs association simulations: characterizing the age bias adjustment maintaining “true” OR with control cohort age progression (best fit power: Δ*T*^*P*^). Common, low-effect-size alleles, showing two SNPs—with the largest and the smallest effect—for each LOD. The confidence interval bars correspond to two-sigma (95%) based on standard error of linear regression fitting. In this plot, rather than using the square of Δ*age*, the best fit power is iteratively discovered, achieving better residual standard error and P-value of the R lm() regression, compared to Figure S6.

**Figure S10.**
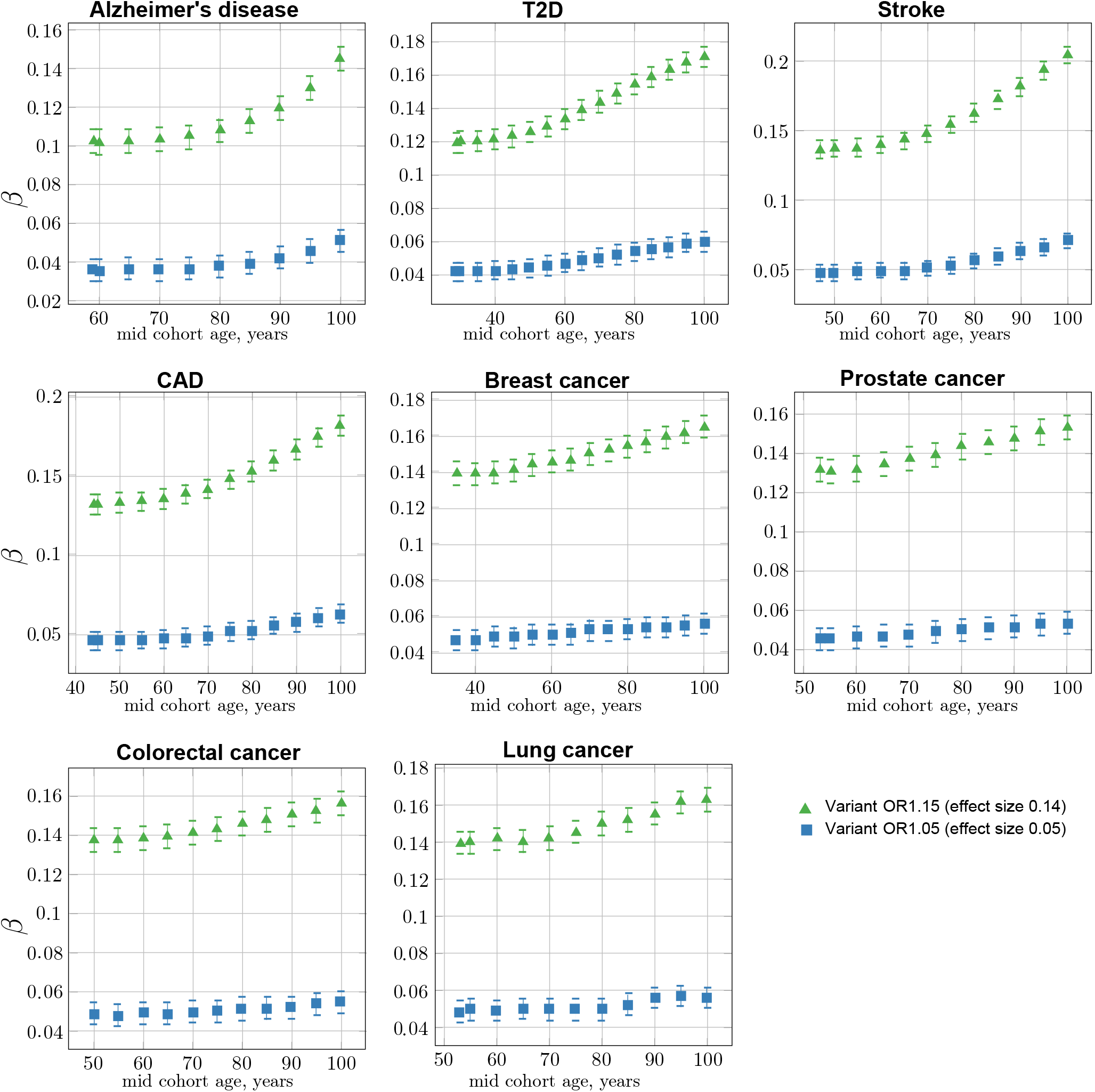
GWASs association simulations: absolute effect size progression with control cohort age increasing against the constant youngest possible case cohort. Common, low-effect-size alleles, showing two SNPs—with the largest and the smallest effect—for each LOD. The confidence interval bars correspond to two-sigma (95%) confidence from the GWASs’ logistic regression association.

**Table S1.**
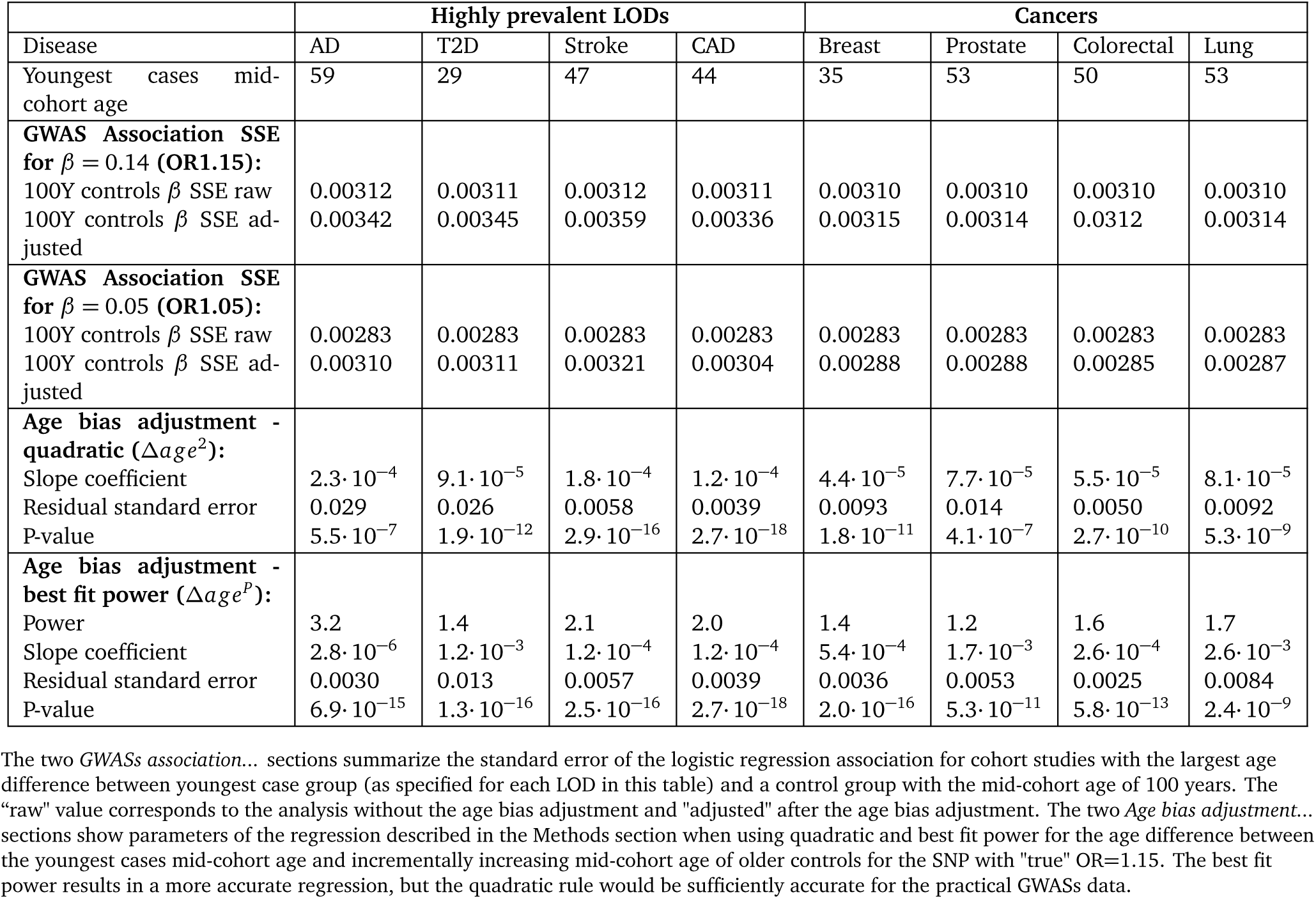
Summary of GWASs association simulations and effect size correction parameters for youngest cases-older controls cohorts.

| Disease | Highly prevalent LODs |  |  |  | Cancers |  |  |  |
| --- | --- | --- | --- | --- | --- | --- | --- | --- |
|  | AD | T2D | Stroke | CAD | Breast | Prostate | Colorectal | Lung |
| Youngest cases mid-cohort age | 59 | 29 | 47 | 44 | 35 | 53 | 50 | 53 |
| **GWAS Association SSE for $\beta = 0.14$ (OR1.15):** |  |  |  |  |  |  |  |  |
| 100Y controls $\beta$ SSE raw | 0.00312 | 0.00311 | 0.00312 | 0.00311 | 0.00310 | 0.00310 | 0.00310 | 0.00310 |
| 100Y controls $\beta$ SSE adjusted | 0.00342 | 0.00345 | 0.00359 | 0.00336 | 0.00315 | 0.00314 | 0.0312 | 0.00314 |
| **GWAS Association SSE for $\beta = 0.05$ (OR1.05):** |  |  |  |  |  |  |  |  |
| 100Y controls $\beta$ SSE raw | 0.00283 | 0.00283 | 0.00283 | 0.00283 | 0.00283 | 0.00283 | 0.00283 | 0.00283 |
| 100Y controls $\beta$ SSE adjusted | 0.00310 | 0.00311 | 0.00321 | 0.00304 | 0.00288 | 0.00288 | 0.00285 | 0.00287 |
| **Age bias adjustment - quadratic ($\Delta age^2$):** |  |  |  |  |  |  |  |  |
| Slope coefficient | $2.3 \cdot 10^{-4}$ | $9.1 \cdot 10^{-5}$ | $1.8 \cdot 10^{-4}$ | $1.2 \cdot 10^{-4}$ | $4.4 \cdot 10^{-5}$ | $7.7 \cdot 10^{-5}$ | $5.5 \cdot 10^{-5}$ | $8.1 \cdot 10^{-5}$ |
| Residual standard error | 0.029 | 0.026 | 0.0058 | 0.0039 | 0.0093 | 0.014 | 0.0050 | 0.0092 |
| P-value | $5.5 \cdot 10^{-7}$ | $1.9 \cdot 10^{-12}$ | $2.9 \cdot 10^{-16}$ | $2.7 \cdot 10^{-18}$ | $1.8 \cdot 10^{-11}$ | $4.1 \cdot 10^{-7}$ | $2.7 \cdot 10^{-10}$ | $5.3 \cdot 10^{-9}$ |
| **Age bias adjustment - best fit power ($\Delta age^p$):** |  |  |  |  |  |  |  |  |
| Power | 3.2 | 1.4 | 2.1 | 2.0 | 1.4 | 1.2 | 1.6 | 1.7 |
| Slope coefficient | $2.8 \cdot 10^{-6}$ | $1.2 \cdot 10^{-3}$ | $1.2 \cdot 10^{-4}$ | $1.2 \cdot 10^{-4}$ | $5.4 \cdot 10^{-4}$ | $1.7 \cdot 10^{-3}$ | $2.6 \cdot 10^{-4}$ | $2.6 \cdot 10^{-3}$ |
| Residual standard error | 0.0030 | 0.013 | 0.0057 | 0.0039 | 0.0036 | 0.0053 | 0.0025 | 0.0084 |
| P-value | $6.9 \cdot 10^{-15}$ | $1.3 \cdot 10^{-16}$ | $2.5 \cdot 10^{-16}$ | $2.7 \cdot 10^{-18}$ | $2.0 \cdot 10^{-16}$ | $5.3 \cdot 10^{-11}$ | $5.8 \cdot 10^{-13}$ | $2.4 \cdot 10^{-9}$ |

The two GWASs association... sections summarize the standard error of the logistic regression association for cohort studies with the largest age difference between youngest case group (as specified for each LOD in this table) and a control group with the mid-cohort age of 100 years. The "raw" value corresponds to the analysis without the age bias adjustment and "adjusted" after the age bias adjustment. The two Age bias adjustment... sections show parameters of the regression described in the Methods section when using quadratic and best fit power for the age difference between the youngest cases mid-cohort age and incrementally increasing mid-cohort age of older controls for the SNP with "true" OR=1.15. The best fit power results in a more accurate regression, but the quadratic rule would be sufficiently accurate for the practical GWASs data.

